# Indomethacin exerts both cyclooxygenase inhibition-dependent and independent mechanisms to enhance chemo-immunotherapy in mice

**DOI:** 10.64898/2026.01.07.698231

**Authors:** Ogacheko D. Okoko, Nada S. Aboelella, Yan Ye, Xin Wang, Samson E. Simon, Md Yeashin Gazi, Victor J. Enriquez Castro, Mercy Kehinde-Ige, Caitlin Brandle, Dongwen Lv, David W. Wolff, Catherine C. Hedrick, Gary Piazza, Huidong Shi, Gang Zhou

**Author notes:** **Correspondence:** Gang Zhou, Ph.D., Georgia Cancer Center, Augusta University, 1120 15^th^ Street, CN-4140, Augusta, GA 30912. Contribute equally (Nada S. Aboelella, Yan Ye, Xin Wang).

## Abstract

Nonsteroidal anti-inflammatory drugs (NSAIDs) primarily act by inhibiting cyclooxygenases (COX1 and COX2), thereby reducing production of the proinflammatory mediator prostaglandin E₂ (PGE₂). Because PGE₂ is a critical driver of cancer progression and tumor immune evasion, this has motivated interest in combining NSAIDs with chemotherapy or immunotherapy for cancer treatment. However, since COX and PGE₂ levels vary across tumor types, it remains unclear whether tumor PGE₂ abundance solely dictates tumor response to NSAID-based therapies. Here, we investigated the therapeutic potential of indomethacin (Indo), a prototypical NSAID, in combination with cyclophosphamide (CTX), a widely used chemotherapeutic agent with immunostimulatory properties. Metronomic administration of Indo significantly enhanced the antitumor efficacy of CTX in multiple murine tumor models exhibiting variable COX2 and PGE₂ levels, including CT26, MC38, 4T1 and A20. The antitumor effects of CTX+Indo required CD8⁺ T cells and T-cell trafficking from tumor-draining lymph nodes and were further potentiated by anti-PD-1 blockade. Single-cell RNA sequencing (scRNA-seq) revealed that responsive CT26 tumors exhibited a reprogrammed tumor immune microenvironment (TIME), marked by increased effector CD8⁺ T-cell infiltration, reduced immunosuppressive myeloid populations, and enhanced interferon signaling in tumor cells. Importantly, Indo retained therapeutic benefit following CTX even in tumors incapable of producing PGE₂, demonstrating a critical contribution of COX-independent mechanisms, particularly inhibition of tumor-intrinsic oncogenic RAS signaling, to the enhanced efficacy of the CTX+Indo combination. Collectively, our results provide strong preclinical rationale for leveraging the COX/PGE_2_ and RAS dual inhibitory capacities of NSAIDs to enhance the efficacy of chemotherapy and immunotherapy.

## INTRODUCTION

Nonsteroidal anti-inflammatory drugs (NSAIDs) are a class of drugs widely used to treat pain, fever, and inflammation(1). These drugs have also shown strong cancer chemopreventive activity in various rodent models and are associated with reduced incidence of several cancers, particularly colorectal cancer, in epidemiological studies (2–7). The primary mechanism of action of NSAIDs involves inhibition of cyclooxygenases (COX-1 and COX-2), which catalyze the conversion of arachidonic acid to prostaglandin E2 (PGE₂). PGE₂ promotes tumor growth, migration, angiogenesis and chemoresistance(8), and plays a key role in immune evasion by expanding myeloid-derived suppressor cells (MDSCs) and T regulatory cells (Treg) while impairing dendritic cell (DC) function and suppressing T cell and natural killer (NK) cell activities(9–16). The accumulation of PGE₂ within the tumor microenvironment is a well-documented mechanism of immune suppression(17–19). Consistent with this, a multitude of preclinical studies have demonstrated that blockade of the COX/PGE₂ axis can suppress tumor growth while enhancing antitumor immunity(20–22). Among these, NSAIDs such as celecoxib and aspirin have shown promise in improving anti-PD-1 therapy in mouse tumor models(23, 24).

Although many anticancer activities of NSAIDs are attributed to COX/PGE₂ inhibition, additional or alternative COX-independent mechanisms have been described(6, 25–28). For example, prescription NSAIDs such as sulindac, celecoxib, or indomethacin can suppress transcription factors such as NFκB, PPARγ, and β-catenin, which regulate inflammation, transformation, and immune tolerance. They also disrupt oncogenic or pro-tumorigenic signaling pathways, including ERK, AKT, mTOR, WNT, and phosphodiesterases (PDEs), leading to growth arrest and apoptosis. Furthermore, some NSAIDs induce oxidative and endoplasmic reticulum (ER) stress in cancer cells, promoting apoptosis, autophagy, and ferroptosis. These diverse mechanisms, which may sensitize tumors to therapy and/or remodel the tumor immune microenvironment (TIME), have driven interest in combining NSAIDs with frontline chemotherapies or immunotherapies(7, 17, 29).

Among the many commonly used chemotherapeutic agents, the alkylating agent cyclophosphamide (CTX) is unique in its robust immune-potentiating properties. CTX can provoke immunogenic cell death (ICD) that triggers antitumor immune responses, and transiently deplete T regulatory cells (Tregs) and myeloid-derived suppressor cells (MDSCs) to alleviate immunosuppression(30–36). These features have established CTX as an important tool for reprogramming the TIME to support immune-based therapies. Clinically, lymphodepleting CTX is routinely used to precondition patients before adoptive T cell therapy, while preclinical studies have paired CTX with cancer vaccines, therapeutic antibodies, and targeted inhibitors to elicit more robust antitumor immunity(37–40).

We previously reported that Indomethacin (Indo), a prototypical NSAID, enhances the efficacy of adoptive T cell therapy by inducing oxidative stress in tumor cells and sensitizing them to T cell-mediated killing via the TRAIL-DR5 axis(41). In that study, we observed that the combination of CTX and Indo exhibited measurable antitumor activity. Although this effect was less pronounced in the absence of donor T cell infusion, it was significantly greater than that of either agent alone in mice bearing large, late-stage tumors. This observation prompted us to hypothesize that CTX and Indo might show stronger efficacy in treating early-stage tumors. Given that CTX is a core component of many frontline chemotherapy regimens(42, 43), including those for hematologic malignancies, breast and ovarian cancers, sarcomas, neuroblastoma, and retinoblastoma, any therapeutic gain achieved by adding Indo at an early stage of treatment could improve efficacy and prolong survival.

In the current study, we evaluated the CTX+Indo regimen across multiple murine tumor models. We found that CTX combined with Indo consistently outperformed either agent alone in suppressing tumor growth and extending survival. These benefits were immune-dependent and correlated with enhanced T cell function, reduced presence of M2-like macrophages and MDSCs in the TIME, along with diminished oncogenic RAS signaling. Mechanistically, our study reveals that both COX-dependent and COX-independent mechanisms operate in parallel to confer the enhanced efficacy exhibited by the CTX+Indo combination. These findings implicate CTX+Indo as a cost-effective immunomodulating regimen with broad utilities in the context of chemo-immunotherapy.

## RESULTS

### Indo enhances the antitumor activity of CTX in multiple mouse tumor models

To evaluate whether combining CTX with Indo improves control of early-stage tumors, BALB/c mice bearing subcutaneous CT26 tumors were treated with CTX or Indo alone, or with the combination, once tumors reached 60-90 mm² (Figure 1A schema). While each monotherapy provided modest and transient tumor growth delay, all mice eventually succumbed to progressive disease within 30 days (Figure 1A). In contrast, the CTX+Indo combination markedly inhibited tumor growth and extended survival, with 75% of mice alive on day 30 and 25% achieving durable complete remission by day 60 (Figure 1A-B). To assess the generalizability of this therapeutic benefit, the combination regimen was tested in two additional tumor models. In both MC38 colon cancer (Figure 1C-D) and 4T1 breast cancer (Figure 1E-F) models, CTX or Indo alone only modestly delayed tumor progression, whereas their combination elicited more robust antitumor activity. Notably, nearly 40% of MC38-bearing mice achieved complete tumor regression following combination therapy. Collectively, these data demonstrate that the CTX+Indo combination produces enhanced antitumor efficacy across multiple syngeneic tumor models and may represent a broadly applicable therapeutic strategy.

**Figure 1.**
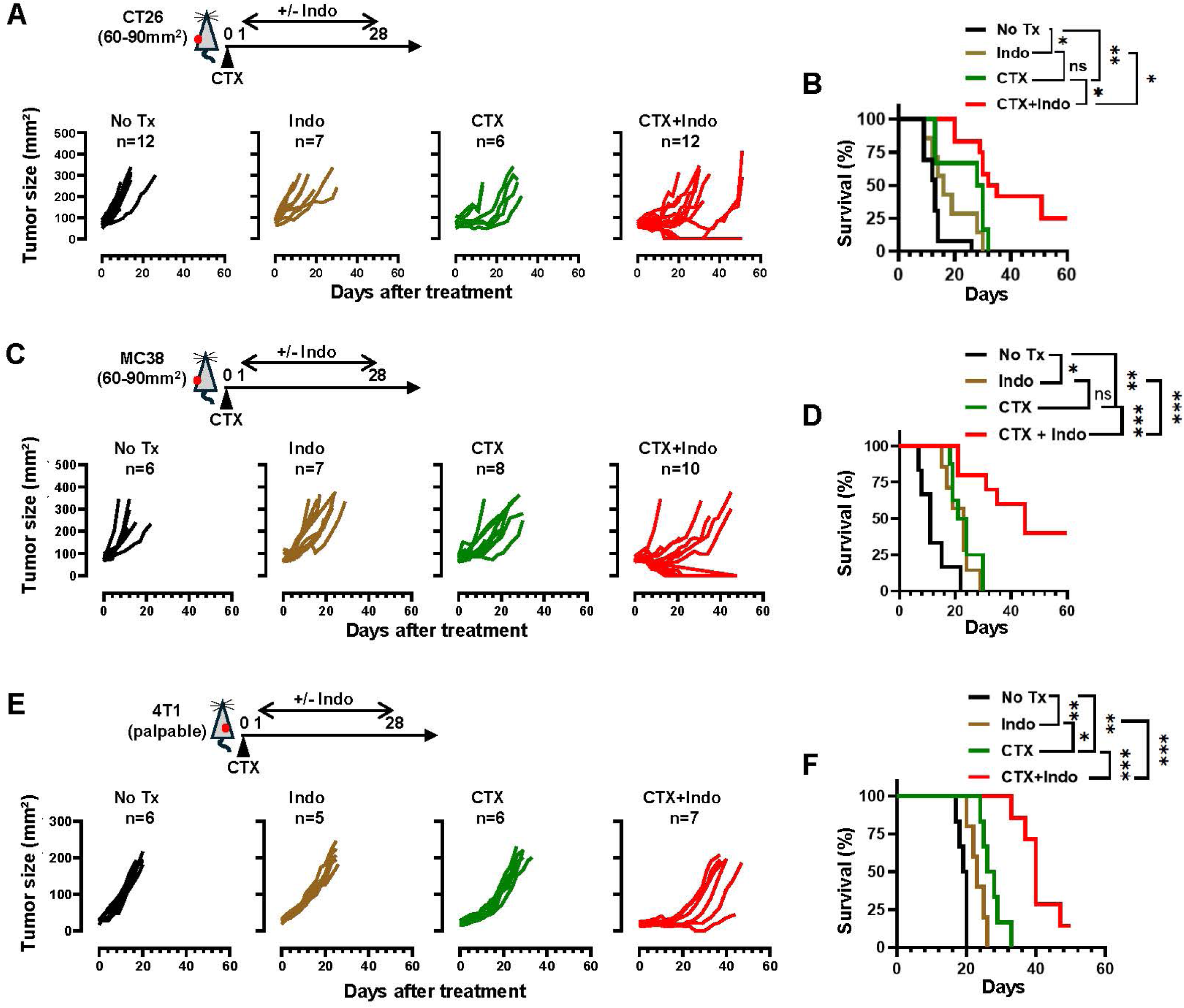
Indo enhances the antitumor activity of CTX in multiple mouse tumor models. (A) CT26 tumor growth curves. The schema depicts the treatment timeline. CT26 cells were implanted s.c. into BALB/c mice. Once tumors reached 60–90 mm², mice were randomized into four groups and treated as indicated. Tumor growth curves are shown, with number of mice indicated. (B) Kaplan–Meier survival analysis of the same cohort. (C) MC38 tumor growth curves. MC38 cells were implanted s.c. into C57BL/6 mice. Mice with established tumors (60–90 mm²) were randomized to the indicated treatment groups. (D) Kaplan–Meier survival analysis. (E) 4T1 tumor growth curves. 4T1 cells were implanted into the fourth mammary fat pad of female BALB/c mice. Mice with palpable tumors (20–30 mm²) were randomized and treated as indicated. (F) Kaplan–Meier survival analysis. Data were pooled from two independent experiments for each tumor model. Statistics: (B, D, F) Log-rank (Mantel–Cox) test. *p < 0.05; **p < 0.01; ***p < 0.001; ****p < 0.0001; ns, not significant.

### The benefits of the CTX and Indo combination treatment require host adaptive immunity

To determine whether the enhanced antitumor effect of CTX+Indo depends on the host immune system, we repeated the tumor treatment experiments in immunodeficient NSG mice (Figure 2A schema). In CT26 tumor-bearing mice, Indo alone did not alter tumor growth kinetics or survival compared with control mice (Figure 2A-B, Indo vs NoTx). Notably, while CTX alone retarded tumor growth and extended survival, the CTX+Indo combination failed to provide additional benefit (Figure 2A-B, CTX+Indo vs CTX), in contrast to the results observed in immunocompetent mice (Figure 1A-B). Similar results were obtained in RAG2-deficient mice (Supplemental Fig. 1), indicating that the host adaptive immune system contributes to the beneficial effects of Indo following CTX chemotherapy. We next examined the role of CD8⁺ T cells, the major cytotoxic effector lymphocytes, in mediating the enhanced antitumor activity of CTX+Indo. To this end, CT26-bearing immunocompetent mice were depleted of CD8⁺ T cells by administration of anti-CD8 antibodies before and during CTX+Indo treatment (Figure 2C schema). Depletion of CD8⁺ T cells abrogated the therapeutic benefits conferred by Indo, rendering the CTX+Indo regimen no more effective than CTX alone in controlling tumor growth or prolonging survival (Figure 2C-D). Together, these findings demonstrate that the therapeutic advantage gained by adding Indo to CTX chemotherapy is dependent on the presence of functional CD8⁺ T cells.

**Figure 2.**
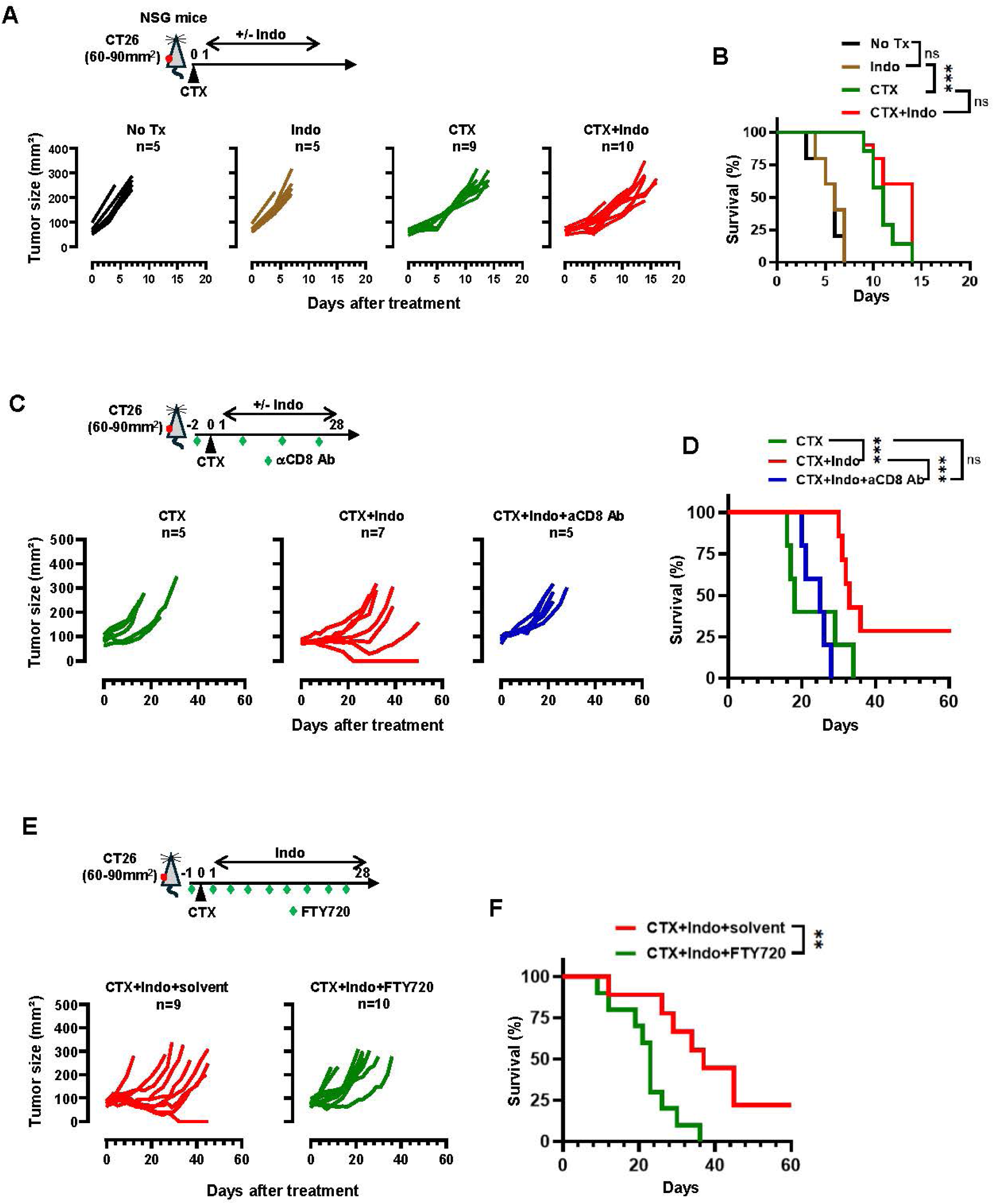
Indo enhances CTX efficacy in an immune-dependent manner. (A) Indo fails to enhance CTX efficacy in immunodeficient NSG mice. The schema depicts the timeline of experimental procedures. NSG mice bearing established CT26 tumors (60–90 mm²) were randomized into four groups and treated as indicated. Tumor growth curves are shown. (B) Kaplan–Meier survival analysis of the same cohort. Data were pooled from two independent experiments. (C) The beneficial effect of Indo requires endogenous CD8⁺ T cells. The schema depicts the timeline of experimental procedures. BALB/c mice bearing established CT26 tumors (60–90 mm²) were treated with CTX+Indo. A subset of mice additionally received i.p. anti-CD8 Ab before and during treatment to deplete endogenous CD8⁺ T cells. Mice treated with CTX alone were included as controls. Tumor growth curves are shown. Mouse survival is summarized in the Kaplan-Meier plot (D). (E) Blockade of T cell trafficking by FTY720 diminishes the efficacy of CTX+Indo. BALB/c mice with established CT26 tumors (60–90 mm²) were treated with CTX+Indo. These mice also received either FTY720 or solvent by i.p. injection three times weekly for 4 weeks, starting one day prior to CTX administration. Tumor growth curves are shown. (F) Kaplan–Meier survival analysis. Data were pooled from two independent experiments. Data were pooled from two independent experiments. Statistics: (B, D, F) Log-rank (Mantel-Cox) test. **, p < 0.01; ***, p < 0.001; ns, not significant.

It is well established that naïve CD8⁺ T cells are primed by dendritic cells (DCs) in the tumor-draining lymph nodes (dLNs), then enter the circulation and migrate to tumors to kill cancer cells. To test whether this process contributes to the antitumor immune response elicited by CTX +Indo treatment, we administered FTY720, a sphingosine-1-phosphate receptor (S1PR) modulator that blocks lymphocyte egress from lymphoid tissues, to CT26-bearing mice during therapy (Figure 2E schema). FTY720 markedly reduced circulating T cells without altering their abundance in secondary lymphoid organs (Supplemental Fig. 2A), confirming effective inhibition of T cell trafficking. Notably, FTY720 treatment attenuated the ability of CTX+Indo to suppress tumor growth and prolong survival (Figure 2E-F). Similar results were observed in the MC38 tumor model (Supplemental Fig. 2B-C). These findings suggest that the therapeutic benefit of CTX+Indo depends on the export of primed CD8⁺ T cells from the dLNs into the circulation and subsequent tumor infiltration to exert cytotoxic activity.

### Single-cell RNA sequencing reveals the impact of Indo on the post-chemotherapy TIME

To characterize changes in the TIME, we performed single-cell RNA sequencing (scRNA-seq) on tumor samples from CT26-bearing mice treated with no therapy, CTX alone, or CTX+Indo (Supplemental Fig. 3A). Since our primary objective was to determine how Indo modulates the TIME in the context of chemotherapy, tumors from the Indo monotherapy group were not included in the scRNA-seq analysis. Because CTX+Indo treatment produced heterogeneous outcomes, ranging from transient tumor growth delay to complete regression (Figure 1), we sought to limit the analysis to tumors that were responsive to the combination regimen. To this end, we adopted a bilateral tumor surgery model previously described by others(44, 45), allowing us to distinguish responsive from nonresponsive tumors. As illustrated in Supplemental Fig. 3B, mice were implanted with tumors on both flanks and treated when both tumors reached 60-90 mm². Seven days later, one tumor was surgically resected and cryopreserved for scRNA-seq, while the contralateral “indicator” tumor was left intact to monitor long-term therapeutic response. Only when the indicator tumor remained stable or regressed over a 3-week treatment period was the matched resected tumor designated as responsive and included in the scRNA-seq analysis. This strategy enabled the collection of treatment-responsive tumors at an early time point when tumor sizes were comparable across groups, minimizing potential confounding effects due to differences in tumor burdens.

t-SNE analysis revealed distinct distributions of cancer cells and immune populations defined by their transcriptomic signatures (Supplemental Fig. 3C-D). Stratifying the t-SNE plots by treatment group highlighted treatment-dependent shifts in TIME cellular composition (Figure 3A). As shown in Figure 3B, cancer cells were predominant in untreated tumors, with a substantial fraction exhibiting active cell-cycling. CTX monotherapy reduced the proportion of cancer cells, particularly the proliferating subset, consistent with the preferential cytotoxicity of chemotherapy toward actively dividing cells. CTX+Indo further decreased the fraction of cancer cells, affecting both proliferating and non-proliferating subsets, in line with enhanced immune-mediated tumor clearance. To more closely examine changes in the immune compartment, we isolated CD45⁺ cells from the scRNA-seq dataset and compared the distribution of immune cells across treatment groups. As shown in Figure 3C, the frequency of CD8⁺ T cells increased following CTX treatment and rose even further in tumors responsive to CTX+Indo. Three macrophage subsets, characterized by expression of *Il1b*, *Fabp5*, or *Basp1,* were identified in untreated tumors. The *Il1b*⁺ subset, which has been reported to exhibit pro-inflammatory, M1-like features (46, 47), expanded in tumors treated with CTX or CTX+Indo. In contrast, the fraction of *Fabp5*⁺ macrophages, which have been reported to display anti-inflammatory, M2-like characteristics (48, 49), was reduced following treatment, with the most pronounced decrease observed in CTX+Indo-treated tumors. Moreover, the proportion of tumor-associated neutrophils, which contain granulocytic MDSCs (gMDSCs), increased after CTX treatment, consistent with chemotherapy-induced reactive myelopoiesis as previously reported(50, 51). Notably, the addition of Indo reduced the presence of gMDSCs in the post-chemotherapy tumor microenvironment.

**Figure 3.**
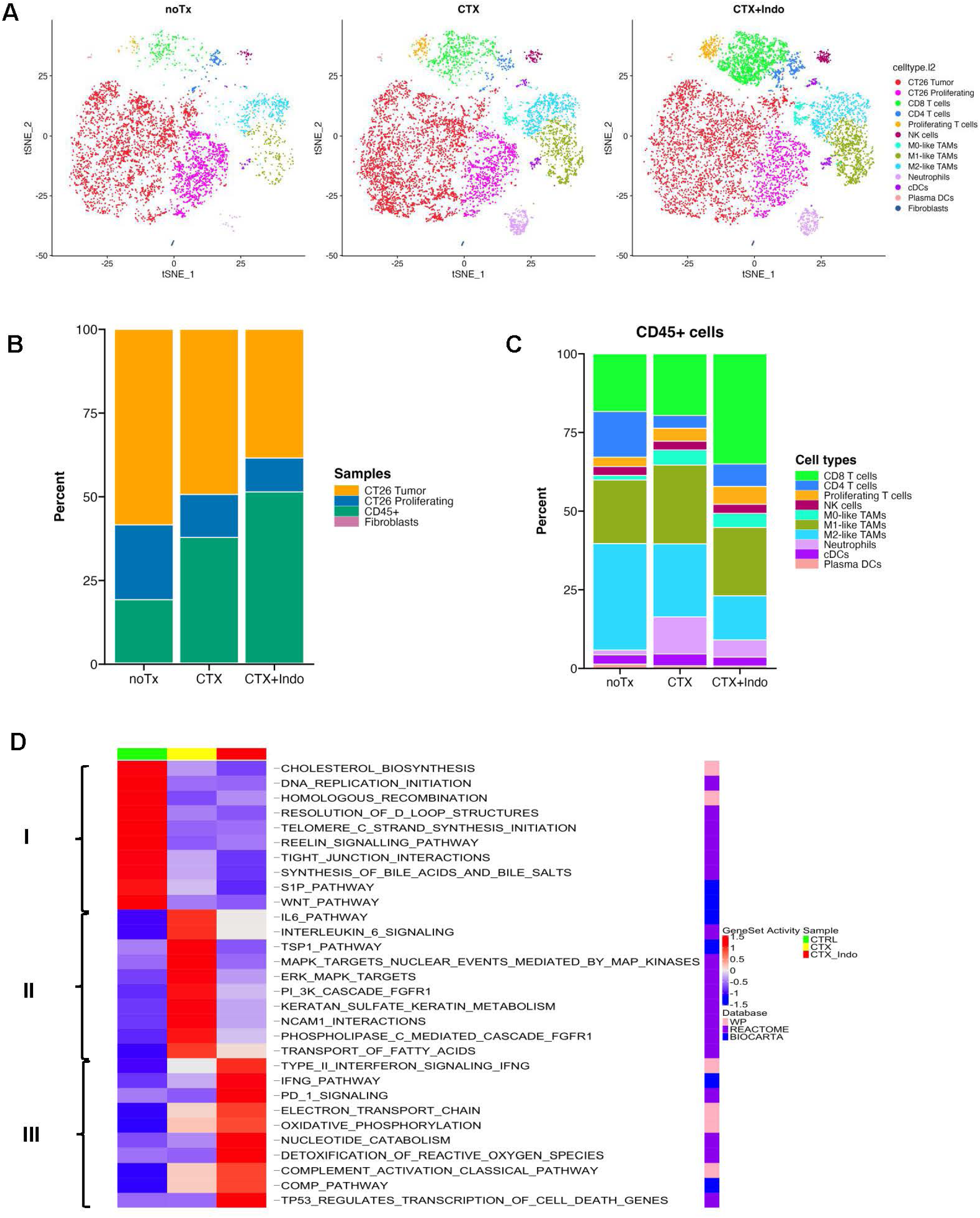
scRNA-seq analysis reveals the impact of Indo on cellular composition and tumor-intrinsic signaling pathways in the post-chemotherapy TIME. BALB/c mice bearing established CT26 tumors (60-90mm^2^) were randomly assigned to three groups to receive no treatment, CTX alone, or CTX+Indo. 7 days after CTX, tumors were harvested, processed and cryopreserved for scRNA-seq analysis. Mice responsive to CTX+Indo treatment (responders) were identified using the bilateral tumor model shown in Supplemental Fig. 3. Only tumors from responders were included in the scRNA-seq analysis. (A) t-SNE plots showing annotated cell populations based on gene expression profiles in the indicated tumor samples. (B) Bar graph summarizing the percentage distribution of tumor cells and immune cells across the analyzed samples. (C) Bar graph summarizing the percentage distribution of different immune cell subsets across the analyzed samples. (D) Heatmap showing the top signaling pathways in tumor cells affected by the indicated treatment conditions. Columns represent samples, and rows represent average pathway AUC scores generated by AUCell package.

We next performed pathway analysis on cancer cells to assess the molecular effects elicited by treatment (Figure 3D). The differentially regulated pathways are clustered into three major groups. Group I comprises pathways enriched in untreated tumors but suppressed by either CTX or CTX+Indo, including cholesterol biosynthesis, WNT signaling, and DNA replication. Group II includes pathways activated by CTX but attenuated upon addition of Indo, including IL-6 and RAS/MAPK (ERK and PI3K) signaling pathways. Group III consists of pathways induced by CTX and further amplified by CTX+Indo, including type II interferon signaling, complement activation, detoxification of reactive oxygen species (ROS), PD-1 signaling, and p53 activation.

Collectively, these data provide insight into how Indo, when combined with CTX chemotherapy, reshapes the TIME by altering immune cell composition and reprogramming cancer cell-intrinsic signaling pathways, shedding light on potential mechanisms through which Indo augments chemotherapy efficacy.

### CTX+Indo augments the efficacy of PD-1 blockade

Because CD8⁺ T cells played a critical role in the antitumor response elicited by CTX+Indo (Figure 2C-D), we next examined their phenotype and function. As shown in Figure 4A, scRNA-seq feature plots revealed increased infiltration of CD8⁺ T cells expressing IFNγ and Granzyme B in tumors treated with CTX+Indo. Notably, these activated CD8⁺ T cells also expressed PD-1, and PD-L1 expression was elevated on macrophages and a subset of cancer cells. To validate the transcriptomic data, we performed flow cytometric analyses on tumors resected 7 days after treatment initiation, using the bilateral tumor model to distinguish between CTX+Indo-responsive (R) and nonresponsive (NR) tumors (Figure 4B). The frequencies of CD8⁺ T cells showed a trend toward increased abundance in CTX-treated and CTX+Indo-responsive tumors relative to untreated controls, although these differences did not reach statistical significance (Figure 4C). Importantly, CD8⁺ TILs from CTX+Indo-responsive tumors exhibited markedly enhanced IFNγ production, both in the percentage of IFNγ⁺ cells and in mean fluorescence intensity (MFI), whereas this enhancement was lost in nonresponsive tumors (Figure 4D-F). Across all treatment groups, the majority of CD8⁺ TILs (>80%) expressed PD-1 (Figure 4G-H), although CTX+Indo modestly reduced PD-1 expression intensity on a per-cell basis (Figure 4I). Together, these observations suggest that CTX+Indo induces tumor-reactive CD8⁺ T cells that remain susceptible to PD-1-mediated dysfunction, raising the possibility that adding PD-1 blockade may further potentiate the therapeutic effect. To test this hypothesis, mice bearing established CT26 tumors were treated with CTX, Indo, or anti-PD-1 antibody, either as monotherapies or in various combinations (Figure 4J schema). Notably, the triple therapy (CTX+Indo+anti-PD-1) produced the most robust antitumor effect, resulting in the highest proportion of durable tumor remissions and prolonged survival compared with all other treatment groups (Figure 4J-K).

**Figure 4.**
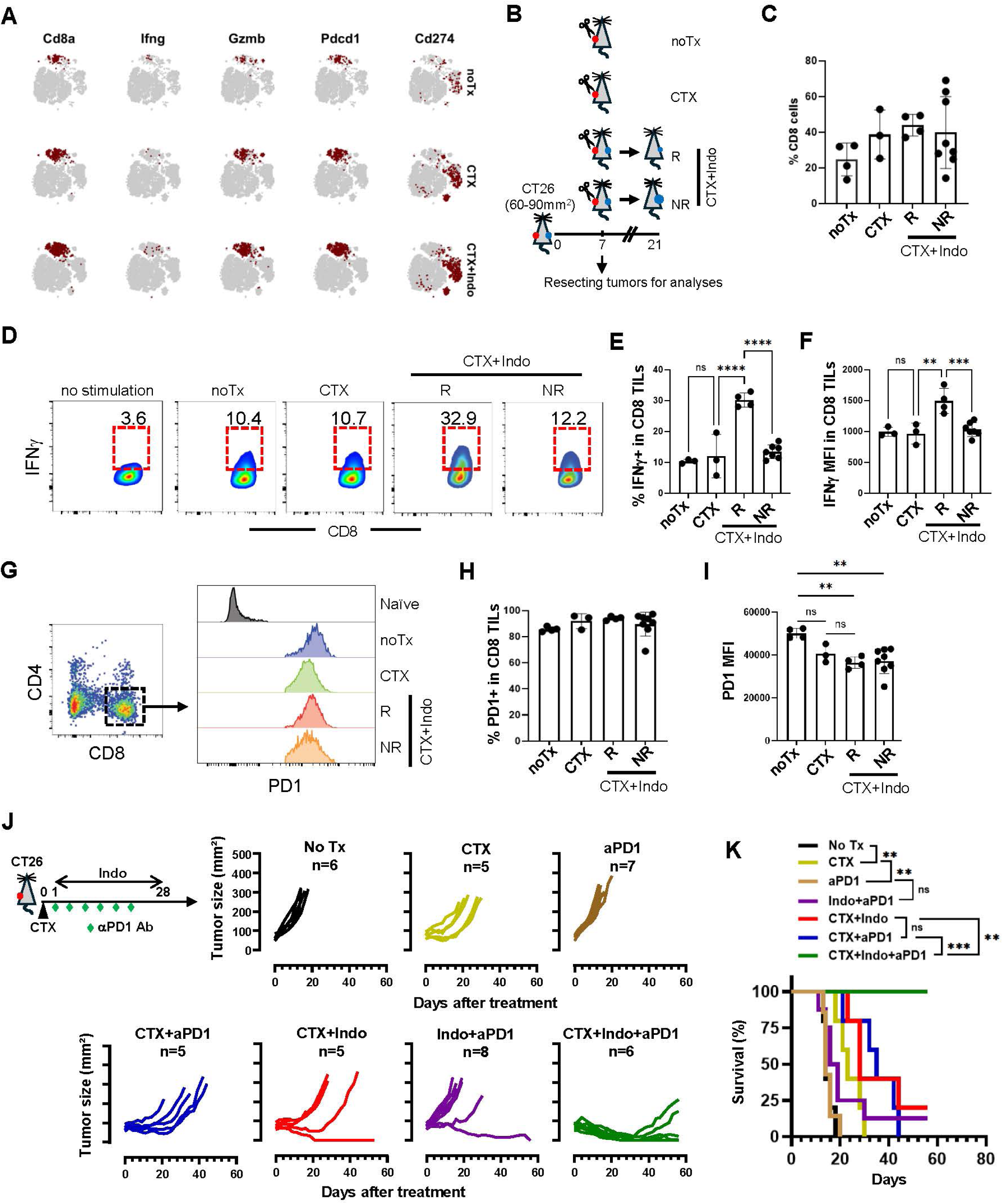
CTX+Indo can augment the efficacy of aPD1 therapy. (A) scRNA-seq feature plots showing the expression patterns of selected genes under the indicated treatment conditions. Data are derived from the scRNA-seq analysis described in Figure 3. (B) Schematic illustrating the timeline of tumor sample collection for flow cytometric analysis. BALB/c mice bearing established CT26 tumors (60-90mm^2^) were randomly assigned to three groups to receive no treatment, CTX alone, or CTX+Indo. 7 days after CTX, tumors were harvested and processed into single cell suspensions for flow cytometric analysis. A bilateral tumor model was used for the CTX+Indo treatment group. Mice responsive (R) or non-responsive (NR) to CTX+Indo treatment were determined based on the outcome of the indicator tumors by day 20, and the corresponding tumor samples analyzed at day 7 were retrospectively categorized. (C) Bar graph summarizing the percentages of CD8+ TILs within lymphocytes under the indicated conditions, shown as mean ± SEM from at least 3 samples per condition. Differences between groups did not reach statistical significance by one-way ANOVA. (D) Representative dot plots showing IFNγ production by CD8+ TILs measured by cytokine intracellular stain after 4-hour stimulation with PMA/inomycin. Numbers denote the percentage of IFNγ⁺ cells within the gated (red rectangles) CD8⁺ T cell population. An unstimulated sample was included to aid in setting the gating strategy. The percentages of IFNγ⁺ CD8⁺ TILs (mean ± SEM) are summarized in (E), and the mean fluorescence intensities (MFIs) of IFNγ in CD8⁺ TILs (mean ± SEM) are summarized in (F). (G) Representative FACS data showing the expression of PD1 in CD8^+^ TILs under the indicated conditions. The percentages of PD1⁺ CD8⁺ TILs (mean ± SEM) are summarized in (H), and the MFIs of PD1 in CD8⁺ TILs (mean ± SEM) are summarized in (I). (J) CTX+Indo augments the efficacy of aPD1 treatment in the CT26 tumor model. The schema depicts the timeline of the experimental procedures. Mice with established CT26 tumors (60-90mm^2^) were randomly assigned to groups to receive the indicated treatments. Tumor growth curves are shown, with the number of mice per group indicated. Mouse survival is summarized in the Kaplan-Meier plot (K). Statistics: (C, E, F, H, I) one-way ANOVA; (K) Log-rank (Mantel-Cox) test. **, p < 0.01; ***, p < 0.001; ns, not significant.

### CTX+Indo mediates antitumor effects through both COX-dependent and COX-independent mechanisms

Numerous studies have established the COX-PGE₂ axis as a key driver of tumor progression and immune evasion, and inhibition of this pathway has been shown to confer antitumor benefits(20–24). Because PGE₂ abundance varies across tumor types(19), we sought to determine whether the therapeutic benefit of CTX+Indo is mediated through COX-PGE₂ inhibition, and whether this dependency is conserved across different tumor models. We first assessed COX-2 expression in a panel of mouse tumor cell lines, including CT26, MC38, 4T1, and A20. Western blot analysis revealed high levels of COX-2 in CT26 cells, followed by lower expression in 4T1 and MC38 cells, whereas COX-2 was undetectable in A20 cells (Figure 5A). As expected, COX-1 was constitutively expressed across all tumor cell lines (data not shown). Corresponding with the COX-2 levels, LC/MS analysis of cell culture supernatants showed abundant PGE₂ production by CT26 cells, followed by lower levels in 4T1 and MC38 cultures, while PGE₂ in A20 cultures remained below the detection limit (Figure 5B). It has been reported that PGE₂ can be produced not only by cancer cells but also by the stromal components (52–54). We thus measured PGE₂ levels in tumor tissues from CT26- or A20-bearing mice. PGE₂ was readily detectable in CT26 tumors but negligible in A20 tumors (Figure 5C). Together, these results demonstrate that the COX-PGE₂ axis is highly active in CT26 tumors but essentially absent in A20 tumors, and that cancer cells are the predominant source of PGE₂ in the CT26 tumor model.

**Figure 5.**
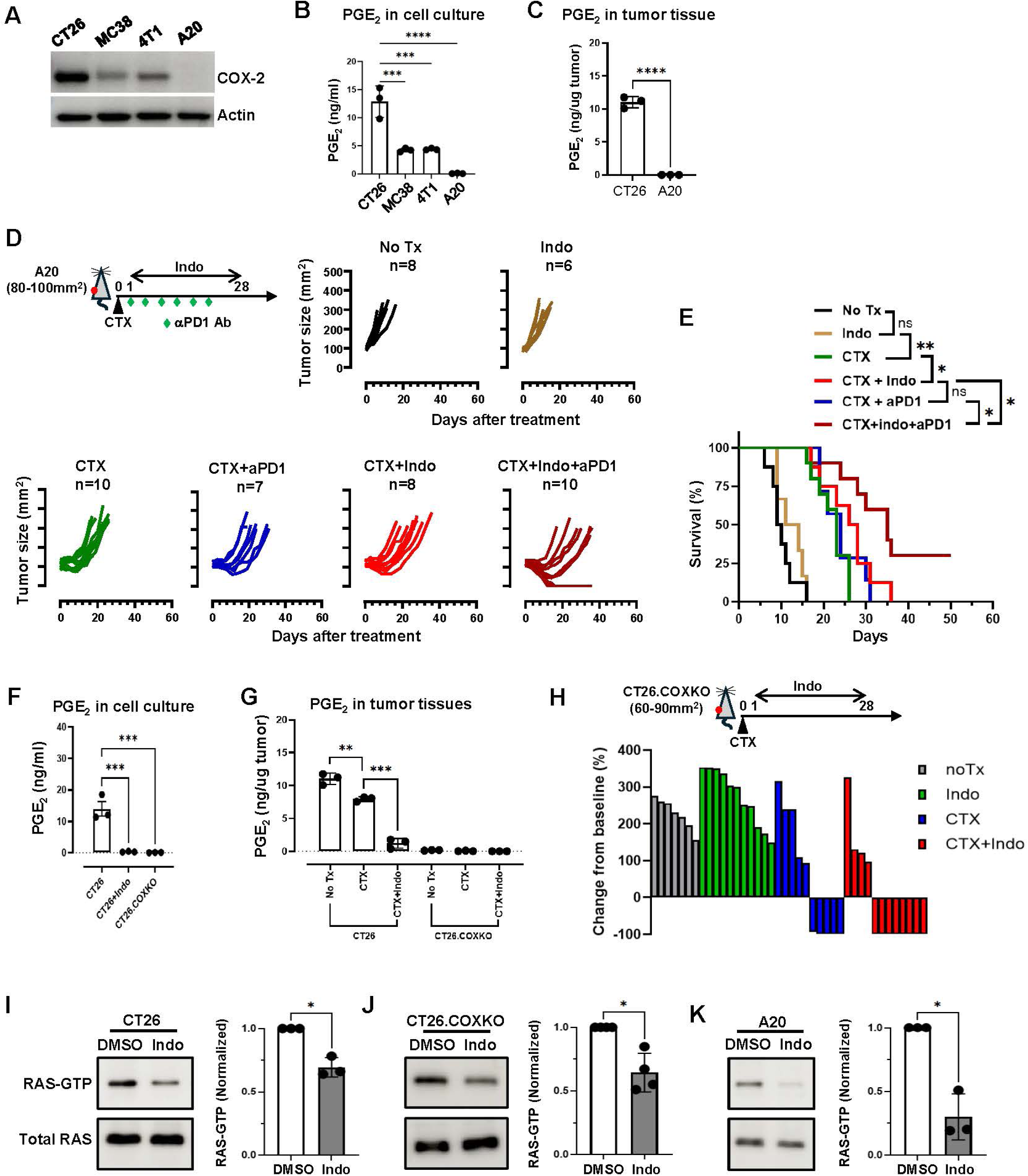
Indo enhances chemo-immunotherapy through both COX-dependent and COX-independent mechanisms. (A) Western blot analysis showing COX-2 expression in the indicated mouse tumor cell lines. β-actin was used as a loading control. (B) PGE₂ concentrations in tumor cell culture supernatants. Tumor cells were seeded in 6-well plates (0.5 × 10⁶ cells per well), and supernatants were collected at confluence. PGE₂ levels were quantified by LC-MS. Data are shown as mean ± SEM of triplicate samples. (C) Quantification of PGE₂ in tumor tissues. BALB/c mice were implanted s.c. with CT26 or A20 tumor cells. Tumor tissues were resected at the sizes of 100-200 mm^2^ and processed for PGE₂ quantification by LC-MS. Results were normalized to tumor tissue weight and shown as mean ± SEM of triplicate samples. (D) CTX+Indo augments the efficacy of aPD1 treatment in the A20 tumor model. The schema depicts the timeline of the experimental procedures. Mice with established A20 tumors (80-100mm^2^) were randomly assigned to groups to receive the indicated treatments. Tumor growth curves are shown, with the number of mice per group indicated. Data shown are pooled from two independent experiments. Mouse survival is summarized in the Kaplan-Meier plot (E). (F) CT26.COXKO cells do not produce measurable PGE₂. Supernatants from CT26.COXKO cell culture were collected for PGE₂ quantification by LC-MS. Cell cultures from untreated or Indo-treated wild-type CT26 cells were included as controls. Results are shown as mean ± SEM of triplicate samples. (G) PGE₂ is not detectable in CT26.COXKO tumor tissues regardless of treatment. BALB/c mice with established wild-type CT26 or CT26.COXKO tumors (60-90mm^2^) were randomly assigned to three groups to receive no treatment, CTX alone, or CTX+Indo. 7 days after CTX, tumors were harvested and processed for PGE₂ quantification by LC-MS. Results were normalized to tumor tissue weight and shown as mean ± SEM of triplicate samples. (H) Waterfall plots summarizing the responses of CT26.COXKO tumor response to the indicated treatments. As depicted in the schema, mice with established CT26.COXKO tumors (60-90mm^2^) were randomly assigned to groups to receive the indicated treatments. Tumor size changes from treatment initiation to endpoint were normalized to the initial tumor sizes for each mouse and used to generate the waterfall plots. (I) Indo reduces the level of RAS-GTP in CT26 cells. CT26 cells were treated with DMSO or Indo (10uM) for 16 hours before being harvested for RAS pull-down assays. Similar assays were conducted for CT26.COXKO cells (J) and A20 cells (K). Representative Western blots for RAS-GTP and total RAS are shown. Normalized RAS-GTP values were calculated as the ratio of RAS-GTP to total RAS in Indo-treated samples divided by the corresponding ratio in DMSO-treated samples. Data are summarized as mean ± SEM from at least three biological replicates per condition. Statistics: (B, F, G, I, J, K), one-way ANOVA; (C), student *t-*test; (E) Log-rank (Mantel-Cox) test. *, p < 0.05; **, p < 0.01; ***, p < 0.001; ****, p < 0.0001; ns, not significant.

We next asked whether the beneficial effects of CTX+Indo observed in COX/PGE₂-active tumor models, i.e. CT26, MC38, and 4T1, could be replicated in the A20 tumor model, which lacks measurable COX2 expression and PGE₂ production. As shown in Figure 5D-E, Indo alone had no effect on A20 tumor growth, and CTX monotherapy produced only a transient delay in tumor progression. CTX+Indo modestly but significantly improved tumor control and survival compared with CTX alone. While CTX combined with anti-PD-1 was not more effective than CTX monotherapy, the triple regimen of CTX+Indo+anti-PD-1 led to more durable tumor control and generated long-term survivors. These results indicate that CTX+Indo retains antitumor activity in the PGE₂-null A20 tumor model. To further validate that the therapeutic benefit of CTX+Indo can occur independently of COX/PGE₂ inhibition, we generated CT26 cells deficient in both COX-1 and COX-2 (CT26.COXKO). Nanopore sequencing confirmed successful gene disruption (Supplemental Fig. 4), and single-cell cloning followed by LC/MS analysis verified that CT26.COXKO cells completely lacked PGE₂ production in vitro (Figure 5F). We then established CT26.COXKO explant clones that can reliably form tumors after inoculation into mice. PGE₂ was confirmed to be absent in CT26.COXKO tumor tissues irrespective of treatment (Figure 5G). As expected, CT26 tumors with intact COX pathways exhibited high levels of PGE₂, which were modestly reduced by CTX treatment and markedly lowered upon addition of Indo, validating the robustness of our PGE₂ quantification. As shown in Figure 5H, Indo alone did not affect CT26.COXKO tumor growth. Notably, CTX monotherapy resulted in complete tumor regression in 50% of CT26.COXKO-bearing mice, indicating that loss of tumor-intrinsic COX-PGE₂ signaling sensitized CT26 tumors to chemotherapy. Moreover, CTX+Indo treatment cured 67% of mice, demonstrating that Indo confers additional therapeutic benefit that is independent of COX inhibition. Together, these findings show that the efficacy of CTX+Indo is not solely reliant on COX/PGE₂ blockade and that additional COX-independent mechanisms contribute to its antitumor activity.

To investigate the potential COX-independent mechanisms involved, we examined our scRNA-seq dataset and observed that CTX+Indo treatment downregulated several oncogenic pathways in CT26 tumor cells, including ERK and PI3K (Figure 3D), which are downstream of the RAS signaling pathway. This observation prompted us to evaluate whether Indo affects tumor-intrinsic RAS signaling, an essential hub that integrates mitogenic cues to support tumor growth and metabolism. RAS pull-down assays showed that Indo significantly reduced RAS-GTP, the active form of RAS, in CT26 cells (Figure 5I), which harbor a KRAS G12D mutation known to sustain high RAS activity(55). Importantly, Indo caused a comparable reduction in RAS-GTP in CT26.COXKO cells (Figure 5J), indicating that this effect is independent of COX inhibition. Indo also decreased RAS-GTP in A20 cells (Figure 5K), which inherently lack detectable COX-2 expression and PGE₂ production, further supporting COX-independent RAS inhibition. Given the key roles of RAS in driving cancer cell proliferation, survival, chemoresistance and immune evasion(56–58), the ability of Indo to suppress RAS signaling provides a plausible COX-independent mechanism that contributes to the enhanced efficacy of the CTX+Indo regimen.

## DISCUSSION

As immunotherapies such as immune checkpoint inhibitors (ICIs) and chimeric antigen receptor-modified T cells (CAR-T) continue to transform cancer care, challenges remain in overcoming primary and acquired resistance(59). In addition, the high cost and specialized infrastructure required for many advanced immunotherapies limit global patient access. Thus, developing low-cost, broadly applicable strategies to enhance the effectiveness of existing therapies is an urgent clinical priority.

Here, we demonstrate that the CTX+Indo regimen exerts multifaceted antitumor effects, including immune enhancement, increased tumor immunogenicity, suppression of oncogenic signaling, and augmentation of PD-1 blockade, through both COX-dependent and COX-independent mechanisms. Although long-term NSAID use is not recommended for cancer prevention or treatment because of risk of gastrointestinal, Kidney, and cardiovascular toxicities(1, 60), the dose of Indo used in our mouse studies (1 mg/kg in mice, ∼0.08 mg/kg human equivalent) is far below standard anti-inflammatory doses recommended for humans (75-150 mg/day). Thus, metronomic administration of low-dose Indo is unlikely to elicit the toxicities associated with chronic high-dose NSAIDs. Given the widespread availability, low cost, and favorable safety profile of both agents at low doses, this combination holds strong translational potential.

NSAIDs have long been recognized for their anticancer activity involving the induction of apoptosis induction of neoplastic cells(61). Beyond their canonical inhibition of COX enzymes and suppression of PGE₂, NSAIDs can regulate redox balance, block oncogenic signaling, impair angiogenesis, and suppress metastasis(6, 25–28). The immunomodulatory effects of NSAIDs, through COX/PGE₂ inhibition, have generated interest in repurposing them as immunotherapy adjuvants, with preclinical studies showing synergy between aspirin or celecoxib and ICIs(23, 24). Our findings extend this concept by demonstrating that CTX+Indo enhances anti-PD1 therapy in both PGE₂-sufficient (CT26) and PGE₂-deficient (A20) tumors. This has important clinical implications given the heterogeneity of PGE₂ abundance across cancers. Many solid tumors (e.g., colon, lung, breast) exhibit elevated PGE₂ levels and may be sensitive to COX inhibition, while malignancies with intrinsically low or undetectable PGE₂ production, such as subsets of non-Hodgkin lymphomas and pancreatic cancers (62–64), may still benefit from NSAID-based combinations through COX-independent mechanisms.

While many NSAIDs exhibit antitumor activity in preclinical models, their clinical benefit in humans has largely been restricted to chemoprevention. Moreover, clinical trials combining NSAIDs with standard chemotherapy in advanced lung or colorectal cancer have thus far reported limited therapeutic efficacy(65–68). In our murine studies, Indo alone had minimal impact on established tumors. However, a single dose of CTX followed by repeated Indo administration significantly delayed tumor progression and improved survival compared with either monotherapy. We postulate that this enhanced therapeutic effect arises from the unique interplay between CTX and Indo, whereby CTX initiates antitumor immune responses, and Indo sustains and amplifies these responses. This notion is supported by our scRNA-seq analyses demonstrating that Indo cooperates with CTX to reshape the TIME (Figure 3), characterized by increased infiltration of CD8⁺ T cells and a concomitant reduction in M2-like macrophages and gMDSCs. In addition, Indo complements CTX by simultaneously enhancing tumor immunogenicity and suppressing tumor-intrinsic oncogenic signaling. For instance, Indo amplifies CTX-induced IFNγ signaling and complement activation in cancer cells, while attenuating CTX-induced ERK, PI3K, and IL-6 signaling pathways, thereby limiting the emergence of acquired chemoresistance. Although CTX is not routinely used in the treatment of lung or gastrointestinal malignancies, it could potentially be incorporated into standard chemotherapy regimens in future clinical trials testing NSAID combination strategies.

A key mechanistic insight from our study is the finding that Indo suppresses tumor-intrinsic RAS activity independently of COX or PGE₂. Although a prior report showed that Indo blocks PGE₂-evoked ERK activation in CT26 cells(69), it remained unclear whether Indo directly modulates endogenous RAS signaling in tumor cells. Using two distinct tumor cell lines, A20 cells with inherent PGE₂-deficiency and CT26.COXKO cells genetically lacking COX enzymes, we demonstrate that Indo decreases RAS-GTP levels irrespective of PGE₂ availability. These findings align with previous reports demonstrating that certain NSAIDs, including Indo, can disrupt the spatiotemporal organization of RAS on the plasma membrane in a COX-independent fashion (70, 71). Because inhibition of oncogenic RAS signaling, via either mutant-selective or pan-RAS inhibitors, can suppress tumor progression and augment antitumor immunity(72, 73), our finding that Indo directly suppresses RAS activity reveals a previously underappreciated COX-independent mechanism that likely contributes to its synergy with CTX. Nevertheless, our results do not exclude the possibility of additional COX-independent mechanisms. For instance, our prior work showed that, in adoptive T cell therapy settings, Indo can sensitize cancer cells to TRAIL-mediated T cell cytotoxicity by inducing oxidative stress and upregulating DR5(41). However, this mechanism does not seem to contribute to the efficacy of CTX+Indo here, as DR5-deficient CT26 tumors responded comparably to wildtype CT26 (data not shown), likely due to lack of TRAIL-expressing cells in the endogenous T cell population. Future studies will be needed to define the specific contribution of RAS inhibition by Indo in chemo-immunotherapy settings.

Despite its improved efficacy, the CTX+Indo regimen has limitations. Complete responses were only observed in a subset of animals, underscoring challenges of tumor heterogeneity and resistance mechanisms. Moreover, the therapeutic benefit was dependent on initial tumor burden, with treatment responsiveness declining once tumors exceeded defined size thresholds (∼100 mm² for CT26 and MC38, 60 mm² for 4T1, and 120 mm² for A20). These observations suggest that CTX+Indo may be most effective when applied at earlier stage of malignancy.

In summary, our study sheds light on the molecular mechanisms by which Indo complements CTX to establish an immunogenic tumor microenvironment. Our results provide a rationale for integrating the CTX+Indo regimen into standard-of-care chemotherapies or immune checkpoint blockade strategies to enhance therapeutic benefit across a broad patient population.

## METHODS and MATERIALS

### Cell lines and culture conditions

A20, 4T1, MC38, and CT26 mouse tumor cell lines were purchased from the American Type Culture Collection (ATCC). All tumor cell lines were cultured in RPMI 1640 (HyClone Laboratories) culture medium supplemented with 10% fetal bovine serum albumin (FBS), 1% penicillin/streptomycin (HyClone Laboratories), 1% non-essential amino acids, 1% glutamine and 0.1mM 2-mercaptoethanol at 37°C in a 5% CO_2_ incubator.

### Generation of CT26.COXKO tumor cell line

COX-1 and COX-2 double knockout CT26 cells (CT26.COXKO) were generated using the Alt-R CRISPR-Cas9 System (IDT) and electroporation following the manufacturer’s protocol. Edited cells were subjected to limiting dilution to obtain single-cell clones. Culture supernatants from candidate clones were screened for PGE₂ production by MS/HPLC, and clones with undetectable PGE₂ were selected for further validation. Targeted gene editing was confirmed by Nanopore sequencing, and loss of COX-1 and COX-2 protein expression was verified by Western blot. The guide RNA sequences and PCR primers used for Nanopore sequencing are listed in the table shown below. The death receptor 5 knockout CT26 cells (CT26.DR5 KO) was generated by similar approach following the procedures described in our previous report(41).

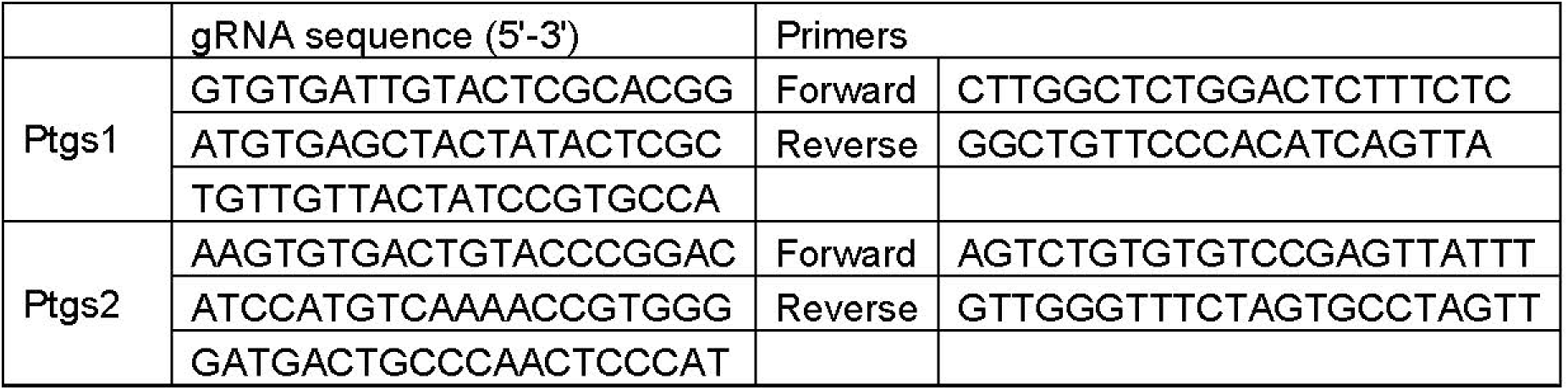

### Antibodies and reagents

Cells were analyzed by flow cytometry using the following fluorochrome-conjugated antibodies CD8-PE/Cy7 (53-6.7), CD4-BV421 (GK1.5), CD45.2-BV421 (104), PD1-PE (29F.1A12), anti– IFN-γ-FITC (XMG1.2). Antibodies were purchased from Biolegend. COX-1 and COX-2 antibodies were purchased from Cayman Chemicals. CTX was purchased from Tokyo Chemical Industry (Tokyo, Japan). Indomethacin was purchased from Sigma-Aldrich. Fingolimod hydrochloride (FTY720) was purchased from MedChemExpress. CD8 depletion antibody (2.43) and anti-PD1 antibody (RMP 1-14) were purchased from BioXCell. GentleMacs tumor dissociation kit was purchased from Miltenyi.

### Experimental mice

BALB/c and C57BL/6J mice (5-10 weeks old) were purchased from Charles River NCI at Frederick. Immunodeficient nonobese diabetic (NOD)-severe combined immunodeficient (Scid) IL-2Rγ-null (NSG) mice were purchased from the Jackson Laboratory. Rag2KO mice lacking the recombination activating gene 2 were (C.129S6(B6)-Rag2^tm1Fwa^N12) purchased from Taconic Biosciences. Both male and female mice were used in study except that only female mice were used for 4T1 breast cancer model. All mice were housed under specific pathogen-free conditions by Laboratory Animal Services of Augusta University. All animal experiments were approved by the Institutional Animal Care and Use Committee of Augusta University.

### Animal tumor models and treatments

A20, CT26 and MC38 tumor cells were subcutaneously implanted to the flanks of mice. 4T1 cells were inoculated to the 4^th^ mammary fat pad of female BALB/c mice. For bilateral tumor studies, CT26 cells were inoculated into both flanks. The following numbers of tumor cells in 50 µl volume were used for each mouse: 2.5×10^6^ for A20, 1×10^6^ for CT26, CT26.COXKO, CT26.DR5KO, MC38 and 4T1. Tumor growth was monitored by caliper measurement every other day. Tumor areas were calculated as the product of two perpendicular diameters in square millimeters (mm^2^). Once tumors reached the sizes specified for each experiment, mice were randomly assigned to treatment groups. CTX was administered intraperitoneally (i.p.) as a single dose of 150 mg/kg when tumors reached the indicated sizes. Indo was administered i.p. at 1 mg/kg, beginning one day after CTX and continued daily for 28 days unless otherwise indicated. Anti-PD-1 antibody was given i.p. at 100 µg per mouse every two days for a total of six doses. To deplete CD8^+^ T cells, anti-CD8 antibody was administered i.p. at 100 µg per mouse once weekly for a total of 4 doses, starting 2 days prior to CTX injection. FTY720 was administered i.p. at 25 µg per mouse three times weekly for three weeks, starting one day prior to CTX administration. Mice were euthanized when tumors reached 20 mm in diameter or if they exhibited signs of terminal illness.

### Sample preparation and flow cytometry analysis

Tumor samples were processed into single-cell suspensions using the GentleMacs tumor dissociation kit according to manufacturer’s protocol. Spleen and blood samples were processed into single-cell suspensions after removing red blood cells with ACK lysing buffer. For surface molecule detection, cells were stained with fluorochrome-conjugated antibodies for 15 minutes at room temperature in the dark. For cytokine intracellular staining, single-cell suspensions were stimulated with the GolgiPlug Leukocyte Activation Cocktail (BD Biosciences) for 4 hours at 37°C 5% CO_2_. Cells were processed for surface staining, followed by intracellular cytokine staining using the Fixation/Permeabilization Solution Kit (BD Biosciences).

### Single cell RNA sequencing library preparation, sequencing, and analysis

Tumor tissues were harvested from mice and processed into single-cell suspensions using the GentleMacs tumor dissociation kit according to the manufacturer’s protocol. Single-cell RNA-seq libraries were generated using the 10x Genomics Chromium platform. The cells were loaded onto the Chromium Controller to encapsulate individual cells in nanoliter-scale droplets with barcoded gel beads. Reverse transcription and cDNA amplification were performed within droplets to label transcripts from each cell uniquely. Libraries were prepared using the Chromium Single Cell 3’ chemistry according to the manufacturer’s protocol and sequenced on an Illumina NovaSeq 6000 sequencer. Raw scRNA-seq data were processed using the 10x Genomics cellranger pipeline (v8) to generate gene-barcode count matrices. Downstream analysis was performed using the Seurat R package (v5.0). Quality control (QC) metrics were applied to filter out low-quality cells based on thresholds for gene count (>300) and mitochondrial gene percentage (<10%). Doublets were removed by the DoubletFinder R package (v2.0.4). Following QC, data were log-normalized and scaled, then subjected to canonical correlation analysis (CCA)-based integration to correct for batch effects and enable joint analysis of multiple samples. Dimensionality reduction was performed using principal component analysis (PCA) and Uniform Manifold Approximation (UMAP). Projection clusters were identified using a shared nearest neighbor (SNN) modularity optimization-based algorithm. Differential gene expression analysis was conducted between clusters using the Seurat FindMarker function, and key findings were visualized using heatmaps, violin plots, and other Seurat-supported visualization functions. Single-cell pathway activity scores were calculated using AUcell v1.26.0. Gene sets were downloaded from MSigDB and the WikiPathways database. The average pathway activities were calculated using Seurat AverageExpression function. Heatmaps were generated using ComplexHeatmaps v2.20.0.

### Nanopore long-read sequencing analysis of the edited genomic locus

Genomic DNA was isolated from genetically edited CT26 COX KO tumor cells and wild-type CT26 tumor cells using DNeasy Blood and Tissue Kit by Qiagen. Polymerase Chain Reaction was done according to manufacturer’s protocol using Q5 Hot Start High Fidelity 2X Master Mix by New England BioLabs. Reverse and forward primers were designed using IDT PrimerQuest Tool. To detect CRISPR/Cas9-induced gene editing events, PCR primers were designed to flank the sgRNA target site, enabling amplification of the edited genomic regions. Purified PCR products were prepared for sequencing using the Oxford Nanopore Ligation Sequencing Kit V14 (SQK-LSK114), following the manufacturer’s protocol. The resulting libraries were sequenced on an Oxford Nanopore MinION device. Basecalling was performed using the Dorado v8 basecaller, generating high-quality sequence reads corresponding to each amplicon. The long-read sequencing data were then visualized and analyzed using the Integrative Genomics Viewer (IGV) to inspect editing outcomes, including indel patterns and structural alterations at the target locus.

### Mass spectrometry for detection of prostaglandin E2 (PGE₂)

To quantify the levels of PGE2 in tumor tissues, mice were sacrificed, and tumors were excised, weighed and cryopreserved in −80°C. Tissue samples were homogenized in pre-chilled 80% methanol (spiked with PGE-d4 as internal standard) together with 0.2ml stainless steel beads (0.1-0.3mm^3^), by high-speed beating using a Blue Bullet spin homogenizer with a speed setting 6 for 3 minutes in cold room (4°C). The homogenized sample was then centrifuged at 16,000g for 10 minutes (4°C). The supernatant was transferred to a new tube and centrifuged under vacuum till dry. The extracted sample was redissolved with 50% methanol for liquid chromatography–mass spectrometry (LC-MS) analysis. To quantify the levels of PGE2 in culture medium (CM), culture supernatants from tumor cell lines were harvested and cryopreserved in −80°C. Pre-chilled methanol was added to CM sample (final methanol concentration is 80%) and then vortexed vigorously for 3 minutes at room temperature, followed by centrifugation at 16,000g for 10 min at 4°C. The supernatant was transferred to a new tube and centrifuged under vacuum till dry. The extracted sample was redissolved with 50% methanol for LC-MS analysis. Separation of PGE₂ was performed with a Phenomenex Kinetex C18 column (100 x 2.1mm, 1.7um) on a Shimadzu Nexera UHPLC system at a flowrate of 0.16ml/min, with a gradient elution from 10% to 60% acetonitrile (with 0.1% formic acid) in 6 minutes. The total analysis time is 15 minutes. The effluent was ionized via ion electrospray in negative mode on a TSQ Quantiva triple-quadrupole mass spectrometry with the following instrument settings: ion spray voltage 2500V, sheath gas 35, ion transfer tube temperature 325°C, aux gas 10, vaporizer temperature 275°C, and FWHM of 0.7 for both Q1/Q3 resolution. The optimal collision energy and RF lens were determined using commercial standards. Transitions 351.2/271.2 and 351.2/333.1 were used as qualifier and quantifier for PGE₂. The integrated peak areas for these transitions were calculated for each sample using Skyline software (version 20.0, University of Washington).

### Western blot analysis

Cells were lysed in ice-cold RIPA buffer supplemented with protease inhibitors (Thermo Scientific Chemicals). Protein concentrations were determined using the BCA assay kit (Pierce), and 20 µg of total protein per sample was resolved on a 10% SDS–PAGE gel (Bio-Rad) along with a molecular weight marker. Proteins were transferred onto a PVDF membrane using a wet transfer system at 200 mA for 2 hours. Membranes were blocked in 5% non-fat milk in TBST for 1 hour at room temperature and incubated overnight at 4 °C with the appropriate primary antibodies. After washing with TBST, membranes were incubated with HRP-conjugated secondary antibodies for 1 hour at room temperature. Protein bands were visualized using SuperSignal West Pico PLUS Chemiluminescent Substrate (Thermo Scientific) and imaged on a ChemiDoc system with automatic exposure settings. Densitometric analysis was performed using ImageJ, and protein levels were normalized to β-actin.

### RAS Pull-down assay

Detection of activated (GTP-bound) RAS protein in cell lysates was performed using an affinity pull-down approach based on the RAS-binding domain (RBD) of the RAS effector kinase RAF1. Tumor cells were treated for 16 hours with either DMSO or Indo (10 µM), then lysed in ice-cold lysis buffer. Lysates were collected after centrifugation at 13,500g for 5 minutes at 4 °C. For RAS-GTP measurement, approximately 500 µg of total protein from each sample was incubated with the Raf-RBD beads (Cytoskeleton Inc., catalog #RF-02). The assay was performed according to the manufacturer’s instructions, with minor modifications to the lysis buffer (50 mM Tris-HCl, pH 7.5; 150 mM NaCl; 1% NP-40; 10% glycerol) supplemented with protease and phosphatase inhibitors. Following incubation and washing, bead-bound RAS-GTP was detected by Western blotting using pan-RAS antibodies (Proteintech, catalog #60309-1-Ig). Densitometric analysis was performed using ImageJ to determine the ratio of RAS-GTP to total RAS across at least 3 biological replicates.

### Statistical analysis

Data were analyzed using GraphPad Prism v9.4.1. Survival curves were compared using the log-rank (Mantel-Cox) test. Comparisons between two groups were performed using unpaired Student’s *t*-tests. For comparisons involving more than two groups, one-way ANOVA followed by Tukey’s multiple-comparison test was used. *P* values < 0.05 were considered statistically significant.

## ACKNOWLEDGMENTS

We thank the Flow and Mass Cytometry Core Facility at Georgia Cancer Center (RRID: SCR_025747) for maintaining and ensuring the smooth operation of the flow cytometers. We acknowledge the support and contribution of the Integrated Genomics Core Shared Resources (RRID: SCR_026483) at the Georgia Cancer Center of Augusta University. We thank Dr. Wenbo Zhi at the Proteomics Core of Augusta University for technical assistance and support.

## Author Contributions

O.O. performed research, analyzed data, and wrote the manuscript; N.S.A., Y.Y., X.W., S.E.S. and M.Y.G. performed experiments; V.C., M.K. and C.B. provided assistance in data analysis; D.L., D.W.W., G.P. and H.S. analyzed data and edited the manuscript. G.Z. conceived the study, designed research, analyzed data and wrote the manuscript.

---

**Supplemental Fig. 1.**
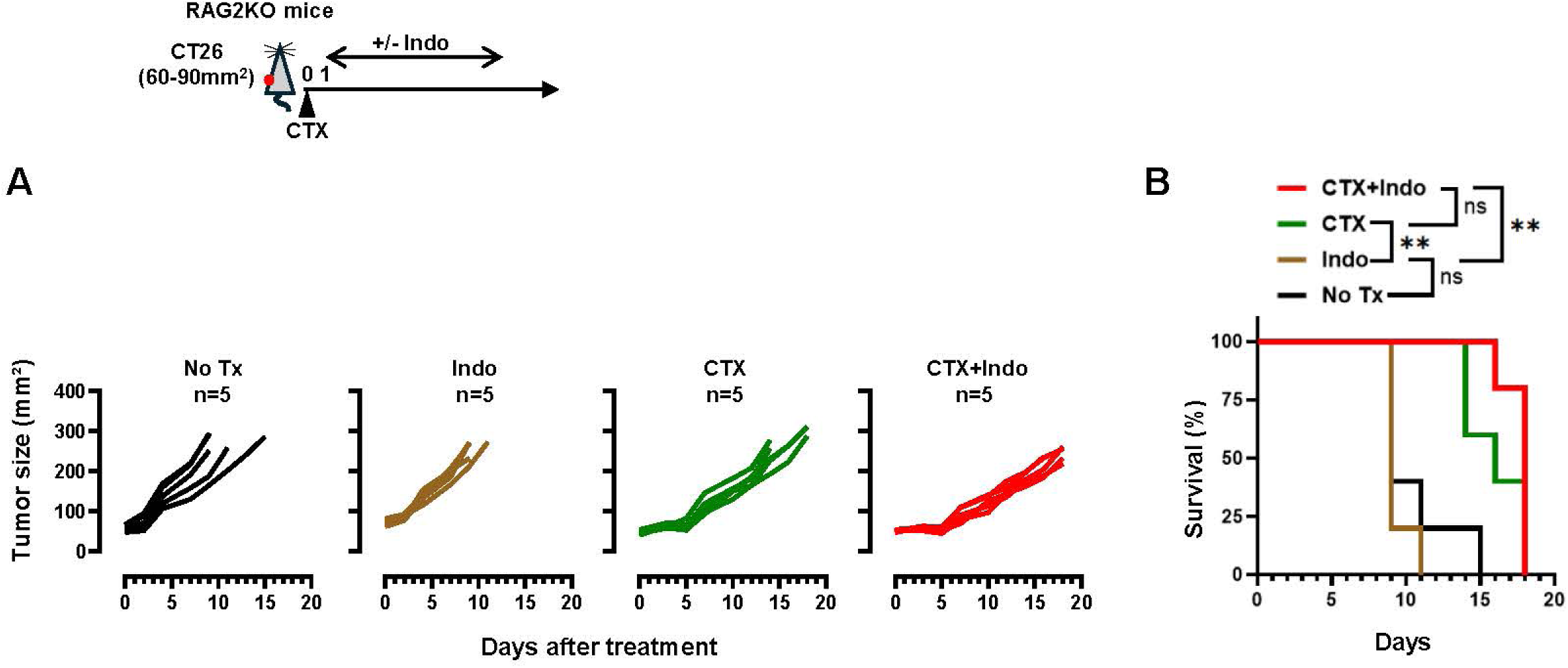
Indo fails to enhance CTX efficacy in immunodeficient RAG2KO mice. The schema depicts the timeline of experimental procedures. RAG2KO mice bearing established CT26 tumors (60–90 mm²) were randomized into four groups and treated as indicated. (A) Tumor growth curves are shown with mice numbers in each group indicated. (B) Kaplan–Meier survival analysis. Statistics: Log-rank (Mantel-Cox) test. **, p < 0.01; ns, not significant.

**Supplemental Fig. 2.**
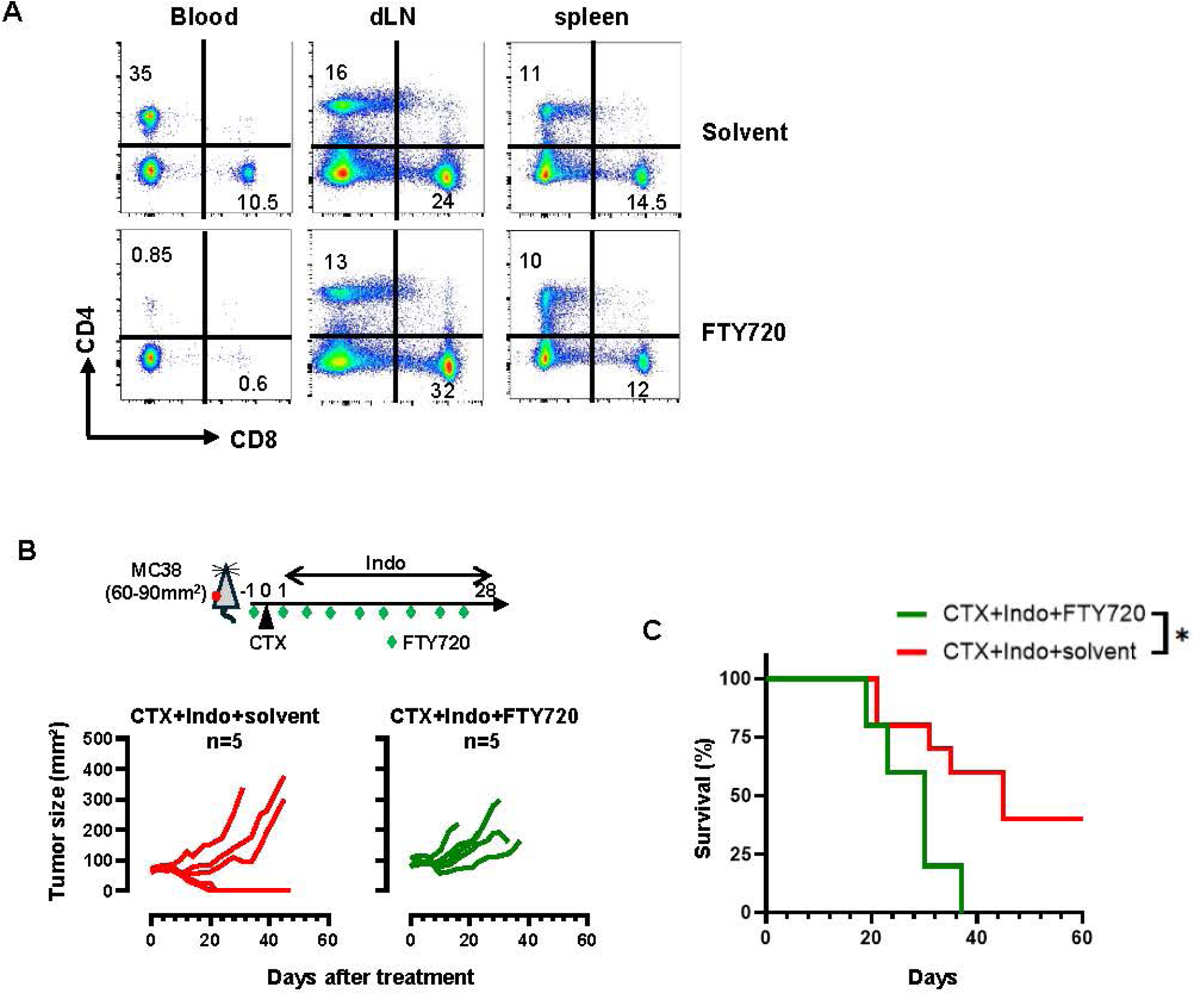
FTY720-mediated inhibition of T cell trafficking diminishes the efficacy of CTX+Indo. (A) FTY720 administration reduces T cells in circulation but does not alter T cell frequency in secondary lymphoid organs. CT26-bearing BALB/c mice were treated as described in Figure 2E. A cohort of mice were euthanized after receiving 4 doses of FTY720 or solvent. The frequencies of CD4+ and CD8+ T cells in blood, draining lymph node and spleen were determined by flow cytometry. Representative dot plots are shown. Numbers in plots denote percent of CD4+ and CD8+ T cells. (B) FTY720 administration diminishes the efficacy of CTX+Indo in MC38 tumor model. Following the same experimental procedures depicted in Figure 2E, C57BL/6 mice with established MC38 tumors (60-90mm^2^) were treated as indicated. Tumor growth curves are shown with mice numbers in each group indicated. Mouse survival is summarized in the Kaplan-Meier plot (C). Statistics: Log-rank (Mantel-Cox) test. *, p < 0.05; ns, not significant.

**Supplemental Fig. 3.**
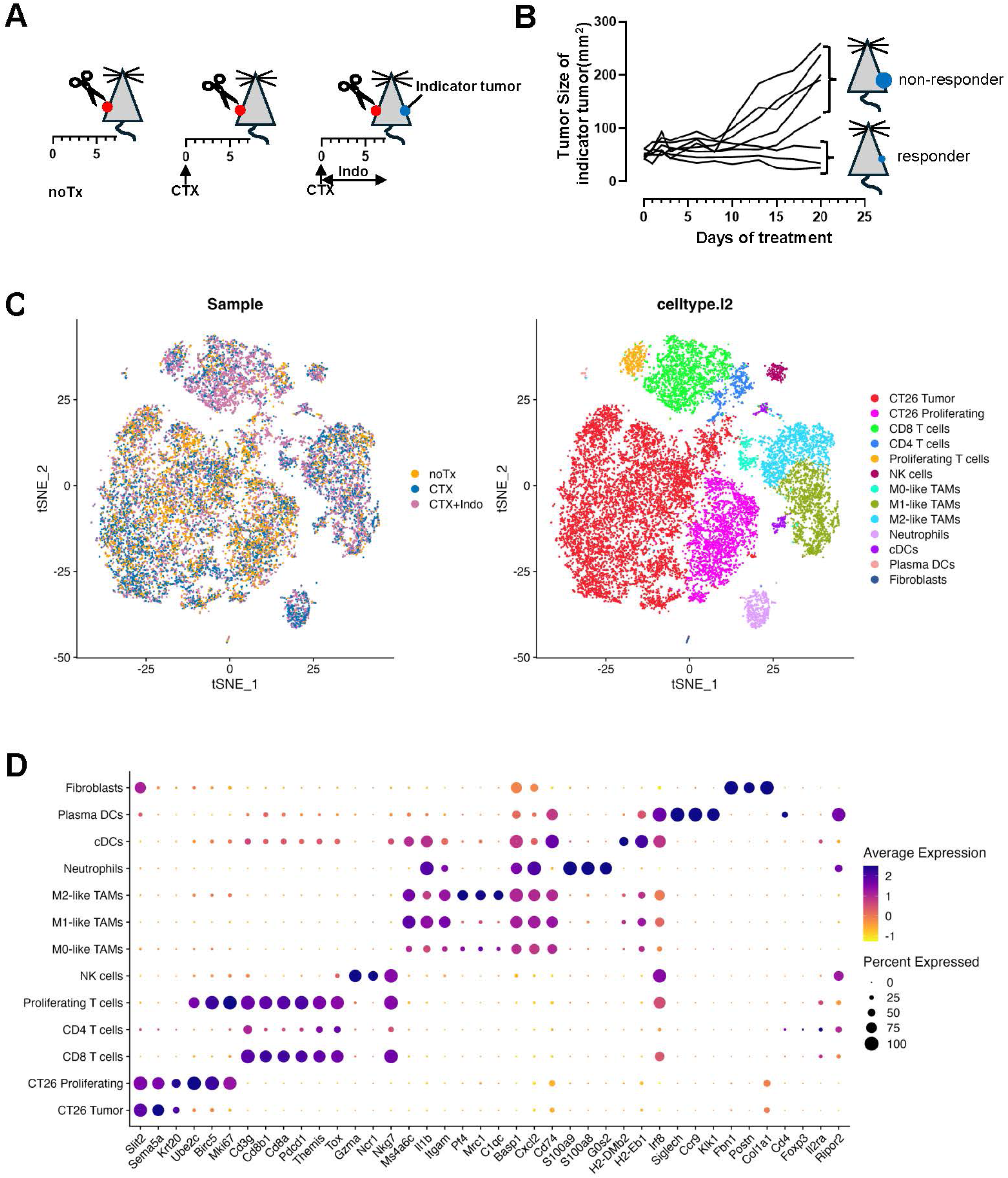
scRNA-seq analysis sample preparation and cell type annotation. (A) Schematic illustrating the timeline of tumor collection under different treatment conditions. A bilateral tumor model was used for the CTX+Indo treatment group. (B) Mice responsive to CTX+Indo treatment (responders) were identified based on regression or stable disease of the indicator tumors by day 20. Only tumors harvested from responder mice, together with tumors from untreated and CTX only-treated mice, were used for subsequent scRNA-seq analysis. (C) t-SNE plots showing annotated cell populations based on gene expression profiles in the indicated tumor samples. (D) Dot plot of cell subtype-specific makers that distinguishes these cell populations. 3 representative genes are shown for each cell subset.

**Supplemental Fig. 4.**
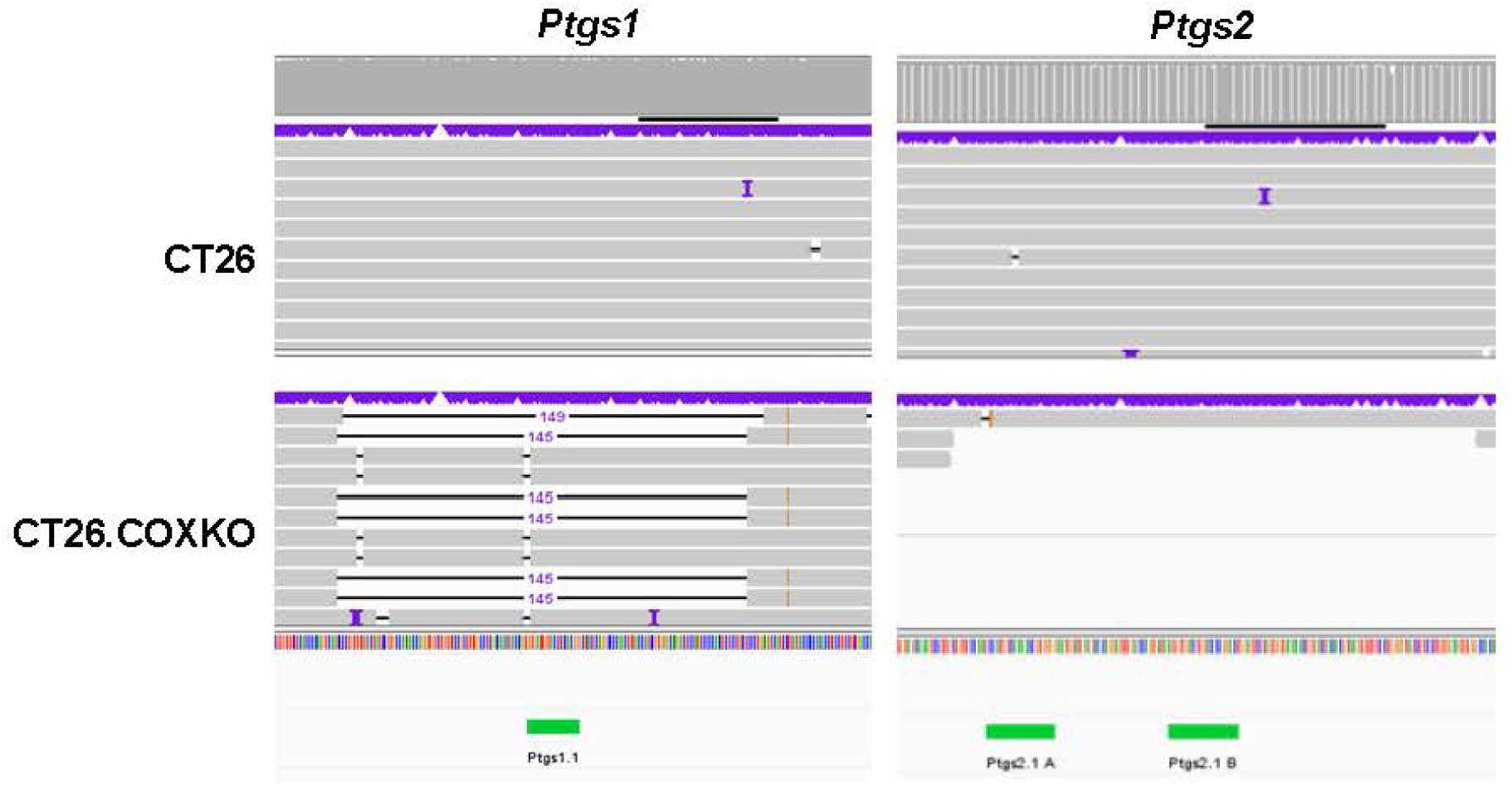
Creation of CT26.COXKO cells. A COX-deficient CT26 cell line was generated by targeted CRISPR/Cas9-mediated gene editing of both *Ptgs1* and *Ptgs2* genes. PCR products spanning the sgRNA target site were amplified and analyzed using long-read sequencing. Images shown are representative IGV browser views of sequencing read alignments illustrating deletions and sequence variants at the CRISPR-edited region. Unmodified CT26 cells were used as a sequencing control.

